# Dynamics of β-cardiac myosin between the super-relaxed and disordered-relaxed states

**DOI:** 10.1101/2024.12.14.628474

**Authors:** Robert C. Cail, Faviolla A. Baez-Cruz, Donald A. Winkelmann, Yale E. Goldman, E. Michael Ostap

## Abstract

The super-relaxed (SRX) state of myosin ATPase activity is critical for striated muscle function, and its dysregulation is linked to cardiomyopathies. It is unclear whether the SRX state exchanges readily with the disordered-relaxed (DRX) state, and whether the SRX state directly corresponds to the folded back interacting-head motif (IHM). Using recombinant β-cardiac heavy meromyosin (HMM) and subfragment 1 (S1), which cannot form the IHM, we show that the SRX and DRX populations are in rapid equilibrium, dependent on myosin head-tail interactions. Some mutations which cause hypertrophic (HCM) or dilated (DCM) cardiomyopathies alter the SRX-DRX equilibrium, but not all mutations. The cardiac myosin inhibitor mavacamten slows nucleotide release by an equal factor for both HMM and S1, thus only indirectly influencing the occupancy time of the SRX state. These findings suggest that purified myosins undergo rapid switching between SRX and DRX states, refining our understanding of cardiomyopathy mechanisms.

## Introduction

Class II myosin paralogs across animalia species form a conserved “off” state in the presence of ATP, in which the two ATPase motor domains of each molecule fold back onto the coiled-coil tail domains, thereby preventing binding to the actin filament and phosphate (P_i_) and ADP release (1). The off-state conformation in striated muscle is termed the interacting-heads motif (IHM) because of inter-head and head-tail interactions responsible for its stabilization (2). Non-muscle and smooth-muscle myosin-II paralogs form a similar state, termed the 10S conformation, that is disengaged upon activation of the muscle through phosphorylation of the regulatory light chains (RLCs) (3, 4). In contrast, skeletal and cardiac muscle myosins do not require light chain phosphorylation for activation. Rather, RLC phosphorylation modulates the partition of molecules into the IHM (5, 6). The two myosin motor head domains in the IHM conformation adopt distinct conformations, with a blocked head that interacts with the myosin backbone and a free head, which binds to the blocked head (7). The conversion between open conformation and IHM/10S conformation is essential to regulating tension maintenance in non-muscle cells, smooth muscle activation, and striated muscle force generation (8–10). Dysregulation of partitioning into the IHM is implicated in many disorders, including hypertrophic cardiomyopathy (HCM) and dilated cardiomyopathy (DCM) (11, 12).

In striated muscle, a biochemically defined population of myosin heads, termed the super-relaxed (SRX) state, saves metabolic energy in the relaxed condition (13, 14). The SRX state was identified in skinned skeletal and cardiac muscle cells using turnover of the fluorescent nucleotide N-Methyanthraniloyl-ATP (mantATP), which is hydrolyzed to mantADP and then released from the myosin heads in two distinct kinetic populations, adequately fitted by exponential decays: a faster-releasing component (at ∼0.05 s^-1^), representing a population termed the disordered-relaxed (DRX) state, and the ∼10-fold slower SRX component (14, 15). X-ray diffraction studies have correlated the SRX and DRX states to myosin heads that are proximal or more distal, respectively, to the backbone of the thick filament (9, 16). The myosin heads that participate in any given contraction are postulated to come from the DRX population. Cardiac SRX heads are thought to stay detached until a regulatory event, such as phosphorylation or length-dependent activation, causes them to shift into the DRX group (9, 14–17).

Are the SRX and IHM states the same? Investigators performing single-turnover kinetic experiments employing purified myosin, either as the dimer-forming heavy meromyosin (HMM) or even the head-only subfragment 1 (S1, which is incapable of entering the IHM state) have reported similar results to skinned striated muscle, with double-exponential fits to mantATP turnover taken to measure the relative proportion and kinetics of SRX and DRX nucleotide turnover (18–21). These results suggest that IHM myosin molecules are in the SRX state. Direct measurement of the kinetics of an SRX-DRX transition, though, has proved difficult (20). X-ray diffraction experiments on myofibrils and FRET studies of purified myosins have undermined the equivalence of SRX with IHM in some conditions, finding evidence for slow-phase nucleotide release that is not correlated to proximal IHM heads (11, 20–22). Moreover, the consistent presence of a measurable slow phase of turnover in S1 is confounding for the head-tail interactions thought to be central to formation of IHM (19–21). A recent publication has reported that a single-exponential decay function is sufficient to model nucleotide release, challenging the reliability of earlier mantATP experiments that were interpreted as the double exponential decay representing the IHM and the open (non-IHM) conformations (23).

The two β-cardiac myosin heads in the IHM conformation are unambiguously in the pre-powerstroke state with ADP.P_i_ in each of the active sites (7). This configuration presumably cannot release P_i_, as the P_i_ release tunnel is blocked by the lever arm. Thus, to undergo P_i_ and nucleotide release, purified myosin molecules seemingly must exit the IHM, with a reaction according to the following scheme:

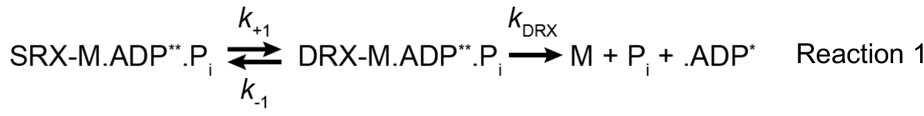

Where M is myosin, ADP** is the high-fluorescence mantADP in the active site and ADP* is the low-fluorescence mantADP in solution.

Whether the IHM (and by implication, the SRX state) is in a rapid equilibrium with the open/DRX state, or whether the two populations are distinct because of a slow transition between the SRX and DRX states, is not yet known (Fig 1A). A double-exponential curve for mantATP turnover suggests that the SRX and DRX populations are separated by a kinetic transition out of the SRX state that is slower than the basal ATPase rate, with this SRX-DRX transition as the rate limiting step for slow nucleotide release. In the case that the equilibrium between the two states is rapid relative to product release, the overall rate of nucleotide release, *k_obs_*, is expected to follow single-exponential kinetics, with an observed rate given by

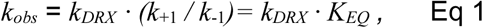

where *k_DRX_* is the elementary product release rate*, k*_+1_ and *k*_-1_ *=* interconversion rates between SRX and DRX (Reaction 1), and *K_EQ_ =* the equilibrium constant for the SRX to DRX isomerization.

**Fig 1:**
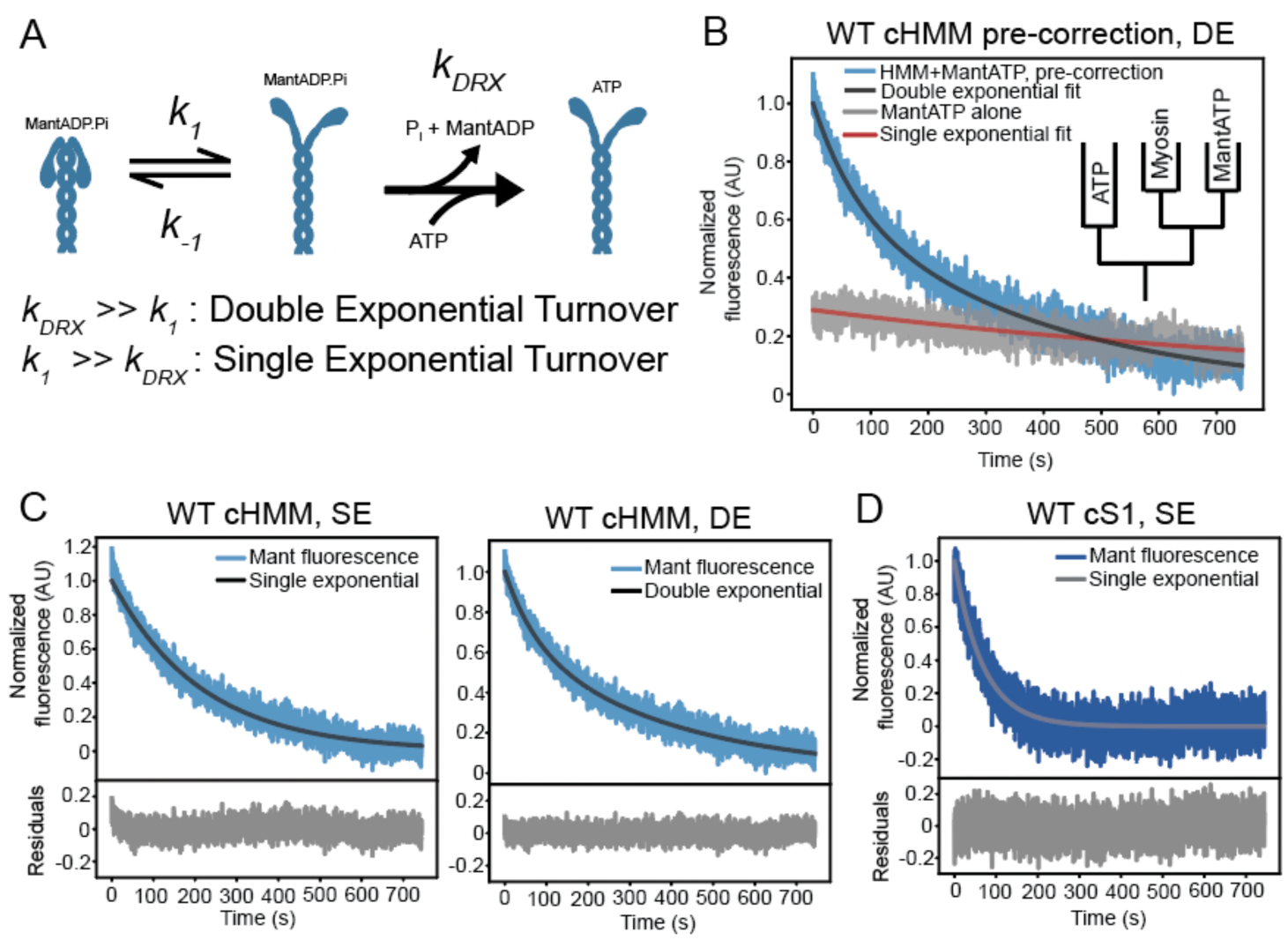
Single vs double exponential equations fitted to mantATP nucleotide turnover data. A) Reaction scheme of SRX-DRX transition followed by release of nucleotide as MantADP. A fast *k*_1_ relative to *k*_DRX_ results in single phase kinetics for nucleotide release rate limited by k_DRX_; if *k*_1_ is substantially slower than *k*_DRX_, the fluorescence decay would be bi-phasic, with the slow phase rate limited by k_1_. B) Single-nucleotide turnover of mantATP from the active site of WT-cHMM (blue) with double-exponential fit, along with a trace of mantATP in the absence of myosin (grey) demonstrating a significant amplitude and rate of mant fluorescence decrease (photobleaching) during kinetic acquisition. Inset: schematic of double-mixing stopped-flow apparatus for experimental conditions employed throughout this paper. C) WT-cHMM single nucleotide turnover after correction for mant photobleaching. Left: double exponential fit with residuals. Right: single exponential fit with residuals. D) WT-cardiac subfragment-1 (-cS1) single nucleotide turnover, corrected for mant photobleaching, with single exponential fit and residuals. SE = single exponential; DE = double exponential.

HCM-causing mutations in *MYH7* (encoding the principal ventricular myosin paralog β-cardiac myosin), *MYLK3* (cardiac-specific regulatory light chain), and *MYBPC3* (cardiac myosin-binding protein C) have been proposed to decrease the number of myosins in SRX, thereby exhibiting an increase in the fast (DRX) amplitude in double-exponential fits and a hyper-contractile state that leads to hypertrophy (24–26). However, it is not clear if disruption of the SRX-DRX distribution is the major driver for disease, as mutations in *MYH7* can also influence other mechanical and biochemical parameters of myosin’s function, such as working stroke and actin affinity (24–28).

In the current work, we employed purified wild-type and HCM mutant cardiac heavy meromyosin (HMM) and subfragment (S1) constructs to investigate 1) the kinetic switching and/or equilibrium state represented by single-nucleotide turnover product release, and 2) how head-tail interactions, myopathy mutations, and small-molecule treatments might alter the SRX/DRX partition.

## Results

### MantATP turnover follows single-exponential kinetics for HMM and S1

We measured the single-nucleotide turnover rate of purified, two-headed, cardiac heavy meromyosin (cHMM) using stopped-flow fluorimetry. In this assay, nucleotide-free myosin is mixed with 1.1-fold excess of mantATP, aged for 10 s to allow nucleotide binding and hydrolysis, and then chased with 1 mM unlabeled ATP to outcompete mant nucleotide as it is released from the myosin active site which decreases the fluorescence intensity (Fig. 1B, inset). Under the experimental conditions in the absence of actin, the rate limiting transition is P_i_ dissociation from the myosin·ADP·P_i_ complex. ADP is released promptly thereafter which generates the observed signal. We employed 295-nm excitation, which excites the mant fluorophore mainly via FRET from Tryptophan 507 near β-cardiac myosin’s nucleotide binding pocket (27).

Single turnover experiments performed with wild-type β-cardiac heavy meromyosin (WT-cHMM) resulted in a biphasic decrease in fluorescence intensity that was well fitted by a double-exponential decaying function, with a slow phase (*k*_s_ = 0.0032 ± 0.0002 s^-1^) contributing a larger amplitude (*A*_s_ = 0.80 ± 0.05) than the fast-phase (*k_f_* = 0.016 ± 0.003 s^-1^; *A*_f_ = 0.20 ± 0.05) (Fig 1B, Fig S1B, Table S1). Because the excitation spectrum of mantATP extends to 295-nm (Fig. S1A), we performed control experiments to determine if changes in fluorescence emission occur independently of myosin kinetics, for instance due to photobleaching. Indeed, over the acquisition window, mantATP fluorescence in the absence of myosin decreased which we attribute to photobleaching (Fig 1B, Fig S1B). The rate of photobleaching, at 0.0017 ± 0.0002 s^-1^, is similar to the slow-phase rate reported above. Because the direct excitation of mantATP at 295 nm is less than the FRET excitation occurring in the presence of myosin, this is expected to be a modest underestimate of the mant photobleaching during myosin single turnover. When the single turnover traces were corrected for photobleaching by subtracting the control traces, transients were well fitted by a single-exponential function with rate of *k*_obs_ = 0.0047 ± 0.0005 s^-1^ (Fig. 1C, Table 1). A two-exponential fit to the corrected transient was not statistically justified according to the Bayesian Information Criterion and the log-likelihood ratio test (Fig. S2, Table S2) (29, 30).

**Table 1:** Single turnover kinetics and equilibrium constant for WT and myopathy mutant HMMs

| Myosin type | Corrected mant-ADP release rate | Papain-digested mant-ADP release rate | $K_{eq}$ |
| --- | --- | --- | --- |
| **WT-cHMM** | $0.0047 \pm 0.0005 \text{ s}^{-1}$ | $0.013 \pm 0.001 \text{ s}^{-1}$ | $0.32 \pm 0.03^a$<br>$0.33 \pm 0.05^b$ |
| **WT-cS1** | $0.0148 \pm 0.0002 \text{ s}^{-1}$ | N/A | N/A |
| **E497D-cHMM** | $0.029 \pm 0.003 \text{ s}^{-1}$ | $0.067 \pm 0.003 \text{ s}^{-1}$ | $0.49 \pm 0.04^b$ |
| **R712L-cHMM** | $0.015 \pm 0.004 \text{ s}^{-1}$ | $0.05 \pm 0.02 \text{ s}^{-1}$ | $0.3 \pm 0.1^b$ |
| **S532P-cHMM** | $0.0035 \pm 0.0003 \text{ s}^{-1}$ | $0.011 \pm 0.002 \text{ s}^{-1}$ | $0.32 \pm 0.05^b$ |
| <sup>a</sup>Calculated assuming $k_{DRX}$ is the rate obtained from the WT-cS1 single-turnover experiment.<br><sup>b</sup>Calculated using $k_{DRX}$ is the rate obtained from the papain-digested cHMM. | | | |

mantATP single turnover assays performed with single-headed wild-type cardiac motor domain (WT-cS1), which is incapable of forming dimers or motor-S2 interactions, resulted in fluorescence transients that were best fit to a single exponential function with a rate of 0.0148 ± 0.0002 s^-1^ after correcting for photobleaching (Fig 1D, Table 1). This rate is >3-fold faster than observed for WT-cHMM (Table 1), and it likely represents the rate of ATP turnover equivalent to the DRX state of myosin (*k*_DRX_).

The faster product release from WT-cS1, coupled with the single exponential decay of fluorescence with WT-cHMM contradicts the model assuming slow escape from SRX (Fig. 1A, ‘slow transition‘ model). Such a slow transition out of the SRX state (<< 0.015 s^-1^) would result in a two-exponential mantATP transient. Rather, the results are compatible with the ‘fast transition’ (>> 0.015 s^-1^) model in Fig. 1A including exchange between SRX and DRX that is rapid relative to the product release step, thus generating a single exponential decay of fluorescence given by Eq. 1 above (Fig. 1A; Table S2) (23).

If we accept that the SRX and DRX states are in equilibrium, we model the SRX-DRX as a rapid equilibrium with the ATPase rate of the DRX state equivalent to that determined for WT-cS1. We calculate the equilibrium constant for the SRX to DRX transition using the observed turnover rate of WT-cHMM (*k*_obs_) as in Eq. 1. *k*_obs_ is taken as the ADP release rate from the DRX state (equivalent to S1 release rate) and *K*_EQ_ is the equilibrium constant of the SRX to DRX transition. This calculation results in an apparent equilibrium constant, *K*_EQ_ = 0.32 ± 0.03 (Table 1). This procedure is conceptually similar to the Long-tail/Short-tail ATPase Ratio (LSAR) assay employed by other groups, in which actin-activated ATPase activity is measured in the presence or absence of the IHM-forming proximal S2 domain (19, 21, 27).

### Intact HMM is required for slowed nucleotide release

We further tested whether IHM interactions in the WT-cHMM molecule are required to slow the single-turnover experiments by performing limited proteolytic digestion of the WT-cHMM with papain, which cleaves myosin at the heavy chain hinge just past the RLC binding IQ motif, releasing S1 (32). Papain digestion was quenched by addition of the irreversible protease inhibitor E-64 (see Methods). SDS-PAGE revealed the proteolytic fragments formed by papain treatment, with predominant bands running at the positions expected for S1, S2, and intact single-headed S1-S2 molecules, none of which are capable of forming IHM (Fig. 2A).

**Fig 2:**
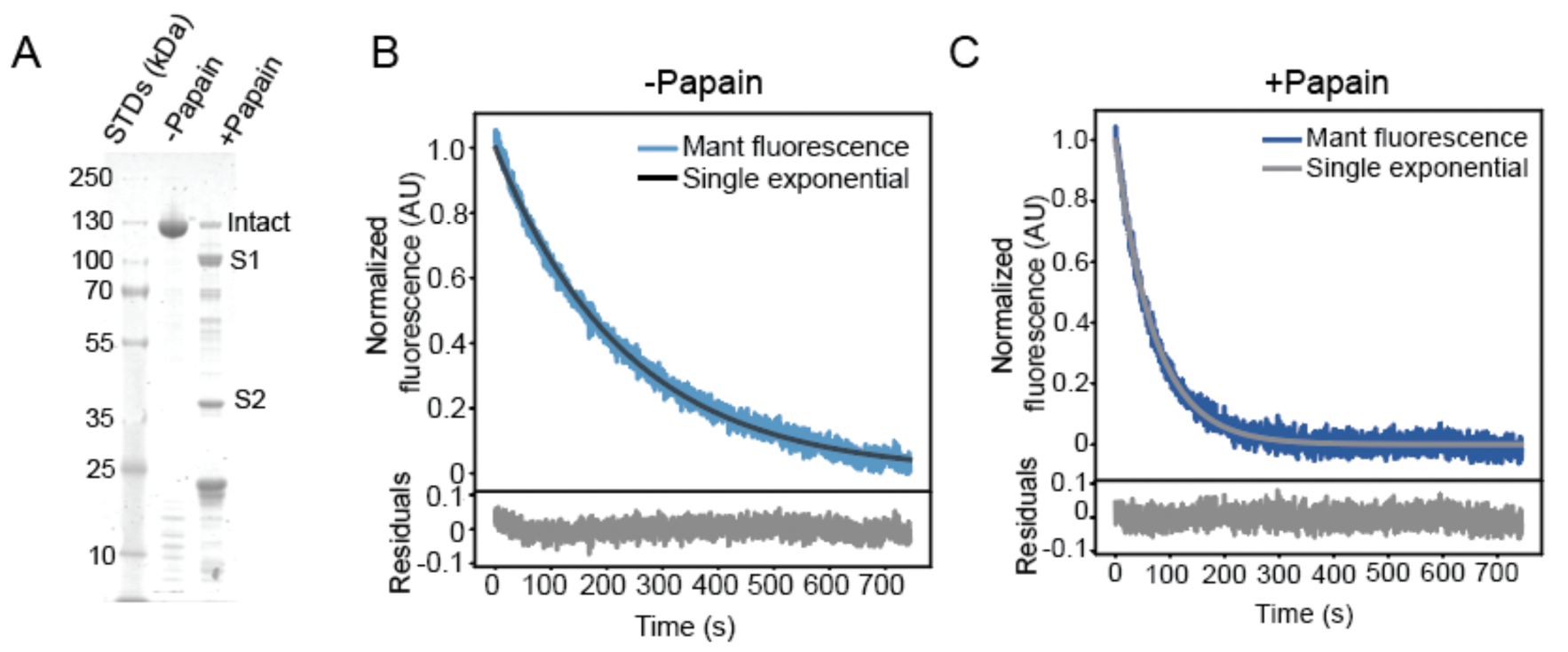
Digestion of WT-cHMM to produce S1 and S2 speeds nucleotide release. A) Coomassie stained SDS-PAGE gel of WT-cHMM demonstrates digestion by papain protease into S1, S2, and other fragments. B) Undigested control samples incubated without the protease have single-turnover kinetics that are well fitted by a single-exponential function, similar to untreated WT-cHMM. C) Upon papain digestion for 5 minutes, the single turnover rate increases significantly to that of WT-cS1, as expected for myosin heads incapable of forming head-tail interactions.

The mantATP turnover rate of papain treated WT-cHMM (0.013 ± 0.001 s^-1^) was similar to WT-cS1, with papain-free control samples (incubated in digestion buffer having identical composition but without papain) retaining the slower single-turnover kinetics (*k*_obs_ = 0.0042 ± 0.0004 s^-1^) observed in untreated samples (Fig 2B-C). The equilibrium constant for digested *vs.* undigested samples, according to Eq. 1, is 0.33 ± 0.05, similar to that of WT-cHMM *vs.* WT-cS1 (Table 1). The presence of a double-exponential mantATP transient was detected at shorter papain incubation times in some experiments, likely indicating the presence of undigested WT-cHMM molecules capable of forming IHM (Fig S3). The results indicate that the IHM-forming interactions between myosin heads and proximal tail domains are essential for the rapid equilibrium that slows nucleotide release in WT-cHMM, supporting the presumption that SRX and IHM are the same state under these experimental conditions.

### Cardiomyopathy mutations can directly alter the SRX/DRX equilibrium, but this is not universal

*MYH7* mutations that cause familial HCM or DCM have been proposed to decrease or increase, respectively, the proportion of IHM/SRX heads, thus altering force production and cooperative thin filament activation (24–29). Published experiments with purified mutant cHMM revealed changes in mantATP transients that have been interpreted to correlate with alteration of the SRX-DRX equilibrium (25, 26). Thus, we investigated the impact of select cardiomyopathy mutations on SRX-DRX equilibrium with the appropriate corrections for photobleaching.

We examined the single-turnover kinetics with cHMM constructs containing HCM mutations (E497D, R712L) and a DCM mutation (S532P). All three mutations demonstrated single-exponential turnover (Fig. S4). E497D-cHMM and R712L-cHMM both exhibited increased the rate of nucleotide release relative to WT-cHMM, with rates of 0.029 ± 0.003 s^-1^ and 0.015 ± 0.004 s^-1^, respectively (Table 1). S532P-cHMM demonstrated little change relative to WT-cHMM (*k*_obs_ = 0.0035 ±0.0003 s^-1^, Table 1). Thus, some but not all of these cardiomyopathy mutations change the basal rate of nucleotide release with S1-S2 interactions intact.

Upon papain digestion, all three mutated myosins had increased rates of nucleotide release. Paired undigested cHMM controls, treated in digestion buffer lacking papain, demonstrated the same product release rate as untreated samples (Fig 3A, Fig 3C, Fig 3E). Upon papain digestion, the rate constants for E497D-cHMM and R712L-cHMM increased to 0.067 ± 0.003 s-1 and 0.05 ± 0.02 s-1, respectively, while S532P-cHMM increased to 0.011 ± 0.002 s-1 (Fig 3B, Fig 3D, Fig 3F, Table 1). With these values, we calculate equilibrium constants for SRX-DRX equilibrium at 0.49 ± 0.04, 0.3 ± 0.1, and 0.32 ± 0.05 for E497D, R712L, and S532P respectively (Table 1). Thus, E497D alters the SRX-DRX equilibrium, but R712L and S532P do not alter it.

**Fig 3:**
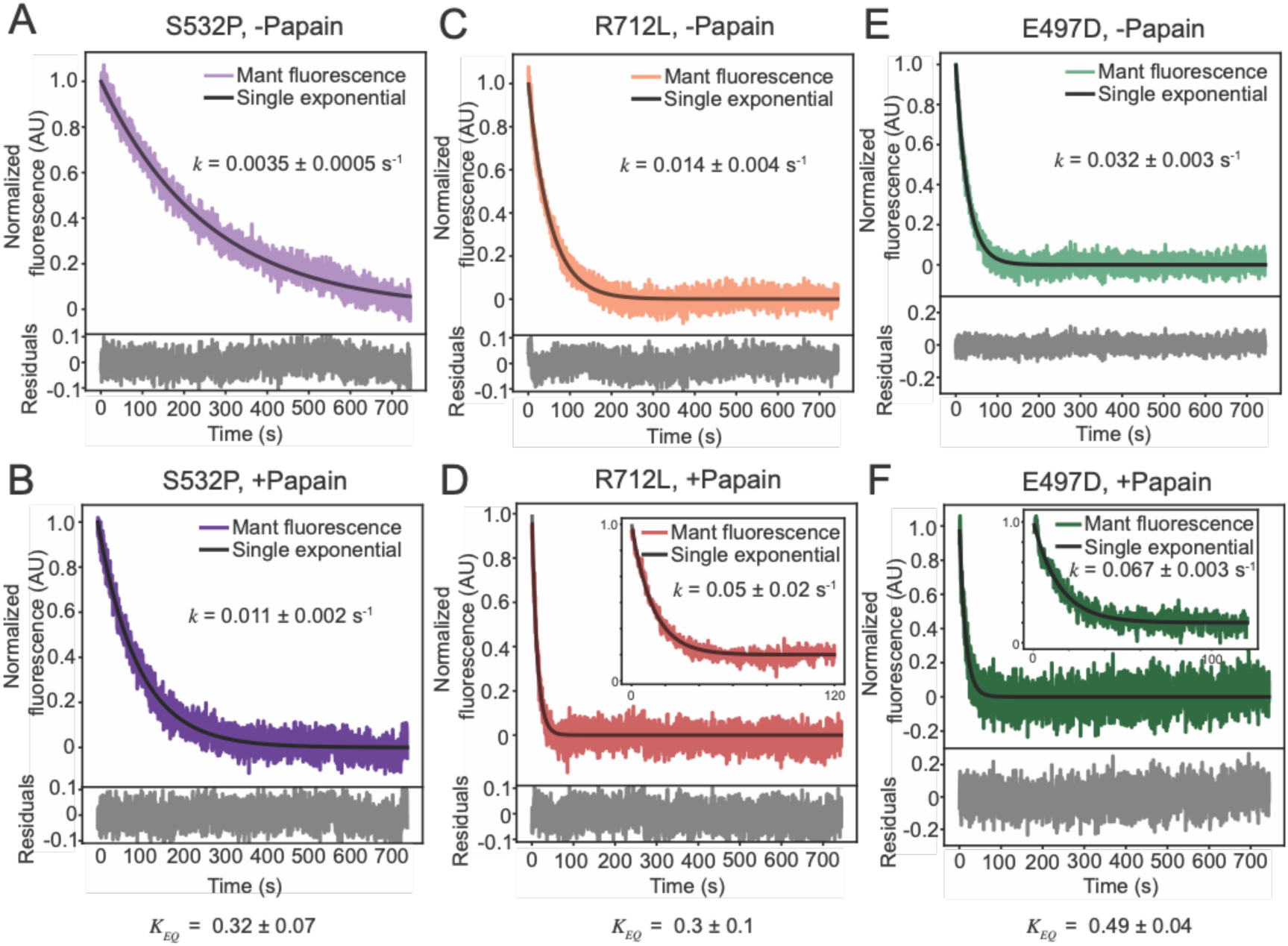
Mutant-cHMM molecules increase single-turnover rate upon proteolytic digestion with papain. A-B) Turnover rate for S532P-cHMM increases from 0.0035 ± 0.0005 s-1 to 0.011 ± 0.002 s-1 upon papain digestion, resulting in a K_EQ_ of 0.32 ± 0.07, similar to WT. C-D) Turnover rate for R712L-cHMM increases from 0.016 ± 0.004 s-1 to 0.05 ± 0.02 s-1 upon papain digestion, resulting in a K_EQ_ of 0.3 ± 0.1, similar to WT. Inset: turnover of papain-digested R712L on a faster time-base to show single-exponential fit for early timepoints. E-F) Turnover rate for E497D-cHMM increases from 0.032 ± 0.003 s-1 to 0.067 ± 0.003 s-1 upon papain digestion, resulting in a K_EQ_ of 0.49 ± 0.04, significantly higher than WT. Inset: turnover of papain-digested E497D on a faster time-base to show single-exponential fit on early timepoints.

### Mavacamten slows nucleotide release without affecting K_EQ_ for SRX-DRX

The drug mavacamten (mava), recently approved to treat obstructive HCM, is well established as an inhibitor of myosin activity in intact muscle cells, skinned muscle fibers, and isolated myosin thick filaments (33–35). It has been proposed to function in part by stabilizing SRX by sequestering heads in the “off” state (21, 35, 36). Mava is also a potent inhibitor of S1 activity in the absence of the S2; thus it may work at multiple levels to inhibit myosin function and decrease thin filament activation (21).

We performed single-nucleotide turnover mantATP experiments with WT-cHMM and WT-S1 constructs to test mava’s effects on the *K*_EQ_ for SRX-DRX. Mava (10 μM) decreased mantATP turnover by cHMM 3.7-fold compared to a DMSO control, as reported previously (Fig. 4A-B, Table 2) (21). A nearly identical 3.9-fold inhibition of WT-S1 was observed (Fig 4C-D, Table 2)). These results indicate that mava does not substantially change the apparent equilibrium constant for the SRX/DRX transition (*K*_EQ_ = 0.38 ± 0.07) (Table 2) under these conditions. Instead, as with some of the myopathy mutations, it appears to alter ATPase kinetics rather than SRX-DRX equilibrium.

**Fig 4:**
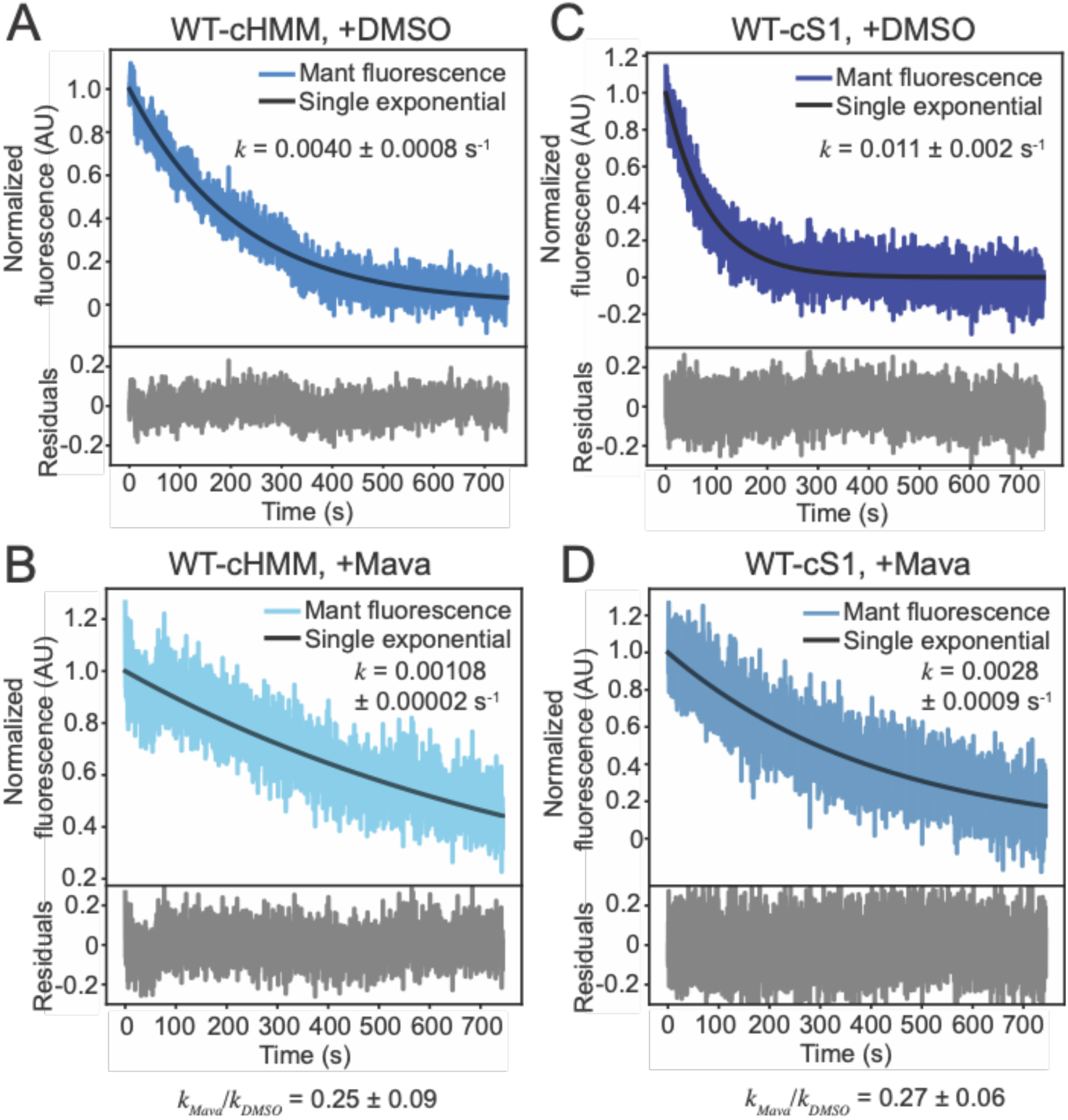
Treatment with mavacamten slows nucleotide turnover by an equal percentage for WT-cHMM and WT-cS1. A-B) WT-cHMM treated with DMSO (A) or 10 μM mava (B) showing a four-fold slowing of nucleotide release in the presence of mava. C-D) WT-cS1 treated with DMSO (C) or 10 μM mava (D), also showing a four-fold slowing of nucleotide release in the presence of mava. The K_EQ_ for the SRX/DRX equilibrium in the presence of mavacamten is 0.38 ± 0.07, statistically indistinguishable from untreated samples.

**Table 2:**
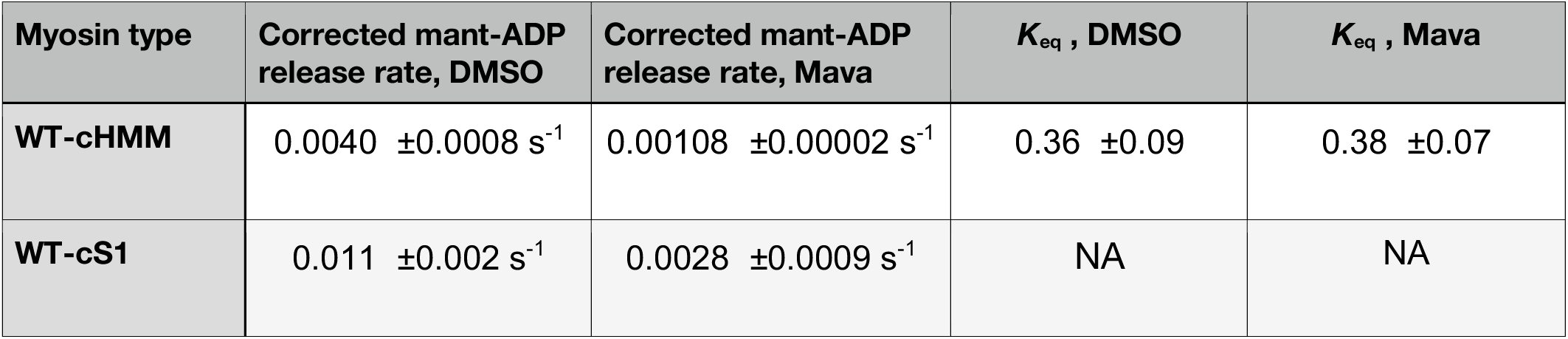
Single turnover in the presence of DMSO or mavacamten

| Myosin type | Corrected mant-ADP<br>release rate, DMSO | Corrected mant-ADP<br>release rate, Mava | $K_{eq}$ , DMSO | $K_{eq}$ , Mava |
| --- | --- | --- | --- | --- |
| WT-cHMM | 0.0040 ±0.0008 s<sup>-1</sup> | 0.00108 ±0.00002 s<sup>-1</sup> | 0.36 ±0.09 | 0.38 ±0.07 |
| WT-cS1 | 0.011 ±0.002 s<sup>-1</sup> | 0.0028 ±0.0009 s<sup>-1</sup> | NA | NA |

## Discussion

We propose that single turnover mantATP transients acquired in the presence of cHMM are best described by an essentially single-exponential process, and that HMM molecules exist in a rapid equilibrium between SRX and DRX. The structure of the IHM state prevents nucleotide exchange, so the dynamic partitioning of motors into the IHM state leads to slower nucleotide turnover, as described in Equation 1. Our findings support previous observations of faster nucleotide release by S1 myosins compared to HMM, but they contradict a recent report of a single-exponential curve for HMM nucleotide release at the same rate as S1 (23). These discrepancies might be attributed to differences in protein preparations (21, 23).

### SRX-DRX in the heart

It is likely that the kinetics of the SRX-DRX transition of myosin *in situ* are slower than the rapid equilibrium kinetics we observed with purified cHMM. Cryo-ET studies revealed that myosin in the intact cardiac thick filament forms three distinct classes templated by unique head/tail positions and interactions with MyBP-C that are not present in purified cHMM (37–39). Therefore, in cardiac muscle, there may exist a kinetically sequestered state that is not attained with cHMM. Indeed, the initial studies that identified a slow turnover population via mantATP were performed with cardiac myocytes (14, 15).

The slow-exchanging SRX myosins act as a motor reservoir ready to be activated in response to requirements for higher cardiac output. If the transition of SRX motors to the DRX state occurs at a rate slower than 0.05 s^-1^, as suggested by double exponential fits of single turnover experiments, then recruitment of the motors would take place on a scale tens or hundreds of seconds which is not compatible with rapid changes in cardiac output. Activation of SRX heads may be the result of regulatory light chain phosphorylation, as observed for non-muscle and smooth myosins (4, 5). However, the Frank-Starling mechanism of stretch-activated muscle output requires a beat-by-beat response, faster than light chain phosphorylation, to increase force production (reviewed in 40). This activation may be the result of the interaction with MyBP-C with myosin, and the presence of the rapidly exchanging SRX-DRX states identified in this study are likely important in this regard.

### Relationship of Mava to the SRX-DRX equilibrium

Mava does not change the SRX-DRX equilibrium of WT-cHMM under the conditions of the present experiments (Fig. 4). However, it does slow nucleotide release from the DRX state. Structural studies reveal that mava stabilizes the Pre-Power-Stroke (PPS) state by interacting with the N-terminal subdomain of the mysosin head and the converter, resulting in a substantial slowing of phosphate release from the active site (34). Importantly, mava does not interact with IHM-forming elements in the region of the motor domain known as the mesa, upper 50 KDa subdomain, or proximal S2 tail. If mava were to increase SRX, we propose that it does so by increasing the proportion of myosins in the PPS that would redistribute into the SRX-DRX equilibrium, rather than changing the SRX-DRX equilibrium directly.

The equilibrium constant for the elementary ATP cleavage step by β-cardiac myosin favors the post-hydrolysis state, and the rate limiting step for nucleotide exchange in the absence of actin is phosphate release (41). This means that most of the motor domains during the ATPase cycle in the absence of actin are in the PPS state. Changes to the equilibrium constant of ATP hydrolysis will change the number of heads in the PPS state, altering the rate of single nucleotide turnover; heads in the PPS state will rapidly form an equilibrium between SRX and DRX conformations. Given this distribution, it is not surprising that the SRX-DRX equilibrium for HMM seems unaffected by mava. These results are largely consistent with previous studies on the effect of mava, which have found at most a modest (4%) difference between IHM and open conformations in the presence of mava, with mava-stabilized heads still available for thin filament interactions upon inotropic stimuli (20, 36).

### The SRX-DRX equilibrium and disease

HCM and DCM are sometimes described as diseases of hyper-contractility and hypo-contractility, respectively, because of their effects on systole, diastole, and left ventricular ejection fraction (25). However, for purified myosins or isolated myofibrils, the effects on force production, kinetic transitions, and tension maintenance are conflicting: some mutations follow the hyper/hypo-contractile phenotype, while others do not (26–29). Destabilizing or stabilizing the IHM is an attractive unifying hypothesis for generating HCM or DCM effects on contractility, irrespective of the mechanochemical changes to individual myosins (25, 29). Our results imply that this model is not universal, but rather the SRX-DRX equilibrium is one of several contributing factors. Cardiomyopathy mutations can change other kinetic parameters such as the nucleotide release rate or the equilibrium constant for ATP hydrolysis ([M.ADP.Pi]/[M.ATP]), which must be considered as possible contributions to disease etiology. For HCM-mutant myosins that reduce the equilibrium constant for ATP hydrolysis, mava would be expected to increase the proportion of post-hydrolysis, PPS myosin molecules (M.ADP.Pi), thus increasing the number of SRX heads irrespective of the SRX/DRX equilibrium constant. Importantly, a parsimonious explanation of mava’s effect is sufficient to explain its utility in treating obstructive HCM: since HCM is a disease of hyper-contractility, inhibition by mava alleviates the symptoms.

## Materials and Methods

### Protein purification

A heavy meromyosin (cHMM) or subfragment-1 (cS1) construct of human β-cardiac myosin (MHY7) was expressed in C2C12 myoblasts and purified as previously described (42). The cHMM or cS1 cDNA was cloned into the pShuttle-IRES-hrGFP-1 vector (Agilent Tech., Santa Clara, CA) and an Ad-Myo-Flag virus was prepared and amplified for expression of cHMM/cS1 protein in C2C12 cells. The cHMM protein has 1146 residues that include residues 1–1138 of the human MYH7 gene and a FLAG tag on the C-terminus (res. 1139–1146). cHMM point mutants were generated by Genewiz (South Plainfield, NJ). The cS1 protein has residues 1-787 of the human MYH7 gene, a 4-residue linker, and residues 5-238 of *A. Victoria* GFP. A FLAG-tagged variant of chimeric protein was prepared by mutating the C-terminal-coding sequence of the GFP domain from DELYK to DYKDHD. Bound light chains are those that are constitutively expressed in the C2C12 cells (MLC1/MLC3 and rLC2).

Confluent C2C12 myoblasts were infected with replication defective recombinant adenovirus (AdcHMM-Flag) at 2.7 × 108 pfu⋅mL−1 in fusion medium (89% DMEM, 10% horse serum, 1% FBS). Expression of recombinant cHMM was monitored by accumulation of co-expressed GFP fluorescence in infected cells. Myocyte differentiation and GFP accumulation were monitored for 216–264 hr after which the cells were harvested. Cells were chilled, media removed, and the cell layer was rinsed with cold PBS. The cell layer was scraped into Triton extraction buffer: 100 mM NaCl, 0.5% Triton X-100, 10 mM Imidazole pH 7.0, 1 mM DTT, 5 mM MgATP, and protease inhibitor cocktail (Sigma, St. Louis, MO). The cell suspension was collected in an ice-cold Dounce homogenizer and lysed with 15 strokes of the tight pestle. The cell debris in the whole cell lysate was pelleted by centrifugation at 17,000 x g for 15 min at 4°C. The Triton soluble extract was fractionated by ammonium sulfate precipitation using sequential steps of 0–30% saturation and 30–60% saturation. The cHMM precipitates between 30–60% saturation of ammonium sulfate. The recovered pellet was dissolved in and dialyzed against 10 mM Imidazole, 150 mM NaCl, pH 7.4 for affinity purification of the FLAG-tagged cHMM on M2 mAb-Sepharose beads (Sigma). Bound cHMM was eluted with 0.1 mg⋅mL−1 FLAG peptide (Sigma). Protein was concentrated and buffer exchanged on Amicon Ultracel-10K centrifugal filters (Millipore; Darmstadt, Germany), dialyzed exhaustively into 10 mM MOPS, 100 mM KCl, 1 mM DTT before a final centrifugation at 300,000 x g for 10 min at 4°C. Aliquots were drop frozen in liquid nitrogen and stored in vapor phase at –147°C. HMM and S1 single-nucleotide turnover experiments A stopped-flow apparatus in sequential mode (SX20 Stopped Flow Spectrometer) was used to acquire all transients for single-nucleotide turnover. The dead time of the instrument is < 3 ms with a total 400-μL sample volume. Fluorescence excitation was provided by a 100-W Hg lamp, where MantATP was excited by fluorescence resonance energy transfer from myosin W508 residue, and nucleotide fluorescence was monitored at 295 nm using a 400 nm long-pass filter for wildtype (WT) and mutant HMM samples. For WT S1-GFP, the same 295 nm was monitored using a 430-470 nm bandwidth filter. All the reagent concentrations reported are post-mixing. For single-nucleotide turnover for WT and mutant HMM, and WT S1-GFP, 50 nM myosin heads were pre-incubated with 52.5 nM MantATP in the aging loop for 10 seconds to allow nucleotide binding and hydrolysis, followed by mixing with 1 mM unlabeled ATP. All proteins and nucleotides were dissolved in KMg25 buffer. For proteolytic fragment formation from HMM to S1, 0.5 mg/mL HMM (pre-incubated with 0.1 U/mL of apyrase-VII on ice for 30-60 minutes before use) was incubated in a total of 60 μL KMg25 with 5 mM cysteine and 18.1 μM papain (diluted from 1.09 mM stock, Sigma-Aldrich p3125) for 3 minutes for mutant HMM and 5 minutes for WT HMM at room temperature. The reaction was quenched with addition of 25 μM E-64 (Cayman Chemical, 10007963); in parallel, a sample was prepared identically but without the addition of papain, to test samples and the effect of papain digestion and subsequent E-64 quenching. From the 60 μL reaction, 15 μL was saved for SDS-PAGE gel analysis, and the remaining was used for stopped-flow.

Samples were incubated on ice until single-nucleotide turnover, which was performed as mentioned above for untreated samples. Mavacamten (10 uM) or an equal amount of DMSO was added to all reagents approximately 5 minutes prior to stopped-flow experiments involving these drugs. Stopped-flow data were acquired using Pro Data-SX software, and fitted to exponentials by a nonlinear least-squares curve fitting. Plots were made using custom python scripts. Statistics were computed in python or microsoft excel. All values are reported as mean ± S.D.

## Data availability

Representative traces from each experiment and paired, unedited SDS-PAGE/coomassie gels are presented in this manuscript; all fits and statistics are reported. Raw data is available upon request from R.C.C. and E.M.O.

## Author contributions

Experiments were conceptualized by R.C.C., Y.E.G., and E.M.O. Adenoviruses were generated and proteins purified by D.A.W. Experiments were conducted by F.A.B.C. and R.C.C., and data were analyzed/figures prepared by R.C.C. Statistical tests were conducted by R.C.C. and Y.E.G. Manuscript was written by R.C.C. and E.M.O. and edited by all authors.

## Competing Interests

The authors declare no competing interests.

## Acknowledgements

This work was supported by the Center for Engineering MechanoBiology NSF Science and Technology Center, CMMI: 15-48571 to Y.E.G. and E.M.O.; , National Institutes of Health grants 5R01HL157997-04 to D.A.W., Y. E. G. and E.M.O., R35GM118139 to Y.E.G., R37GM057247 to E.M.O., and 5T32AR053461 to R.C.C. We thank Richard Wike for technical assistance.

## Notes

### Competing Interest Statement

The authors have declared no competing interest.

