## Supplemental Text and Figures for "Dynamics of β-cardiac myosin between the super-relaxed and disordered-relaxed states"

**This PDF contains Supplementary Materials:**

- Supplementary Methods
- Tables S1-S2
- Figures S1-S5

**Supplementary Methods**

Statistical tests were performed to measure how well single and double exponential models fit the fluorescence decay traces upon Mant turnover from the active site. The Bayesian Information Criterion (BIC, 1) was calculated with the following formula:

$$BIC = n * \ln\left(\frac{SSR}{n}\right) + p * \ln(n)$$

Where $n$ is the number of data points, $p$ the number of model parameters (5 for double exponential and 3 for single exponential), and $SSR$ is the sum of squared residuals for the model fit to the data. A lower value is taken to indicate better model performance.

The log-likelihood ratio (2) was calculated with the following formula:

$$LLR = \frac{loglikehood(single)}{loglikelihood(double)} = \frac{\sum\left(\frac{Sr}{\sigma s}\right)^2 + \ln(2 * \pi * \sigma s^2)}{\sum\left(\frac{Dr}{\sigma d}\right)^2 + \ln(2 * \pi * \sigma d^2)}$$

Where $Sr$ is the residuals of the single exponential fit to the data, $Dr$ is the residuals of the double exponential fit to the data, $\sigma s$ is the standard deviation of the single-exponential residuals, and $\sigma d$ is the standard deviation of the double-exponential residuals. Fluorescence noise is assumed to be normally distributed. A value of 1 indicates the models perform equally well in predicting the data.

**Table S1:** Single vs double exponential characteristics of pre- and post-correction WT myosin

| Myosin type | Fast amplitude | Fast rate | Slow amplitude | Slow rate |
|---|---|---|---|---|
| **WT-cHMM pre correction D.E.** | 0.20 ±0.05 | 0.016 ±0.003 $s^{-1}$ | 0.80 ±0.05 | 0.0032 ±0.0002 $s^{-1}$ |
| **WT-cHMM pre correction S.E.** | N/A | 0.0043 ±0.0002 $s^{-1}$ | N/A | N/A |
| **WT-cHMM post correction D.E.** | 0.15 ±0.03 | 0.020 ±0.003 $s^{-1}$ | 0.85 ±0.03 | 0.0040 ±0.0005 $s^{-1}$ |
| **WT-cHMM post correction S.E.** | N/A | 0.0047 ±0.0005 $s^{-1}$ | N/A | N/A |
| **WT-cS1 pre correction D.E.** | 0.6 ±0.1 | 0.016 ±0.002 $s^{-1}$ | 0.4 ±0.1 | 0.0041 ±0.0002 $s^{-1}$ |
| **WT-cS1 pre correction S.E.** | N/A | 0.0094 ±0.0002 $s^{-1}$ | N/A | N/A |
| **WT-cS1 post correction D.E.** | 0.76 ±0.02 | 0.0147 ±0.0002 $s^{-1}$ | 0.24 ±0.02 | 0.0147 ±0.0002 $s^{-1}$ |
| **WT-cS1 post correction S.E.** | N/A | 0.0147 ±0.0002 $s^{-1}$ | N/A | N/A |

**Table S2:** Statistical comparisons of single and double exponentials

| Trace details | Double Exponential BIC | Single Exponential BIC | % Improvement for double | Log-likelihood ratio |
|---|---|---|---|---|
| **WT-cHMM uncorrected** | -49946 | -42480 | 18 | 0.79 |
| **WT-cHMM corrected** | -43241 | -40451 | 6 | 0.95 |
| **WT-cS1 uncorrected** | -38390 | -31320 | 24 | 0.7 |
| **WT-cS1 corrected** | -25261 | -25245 | 0.1 | 0.99 |

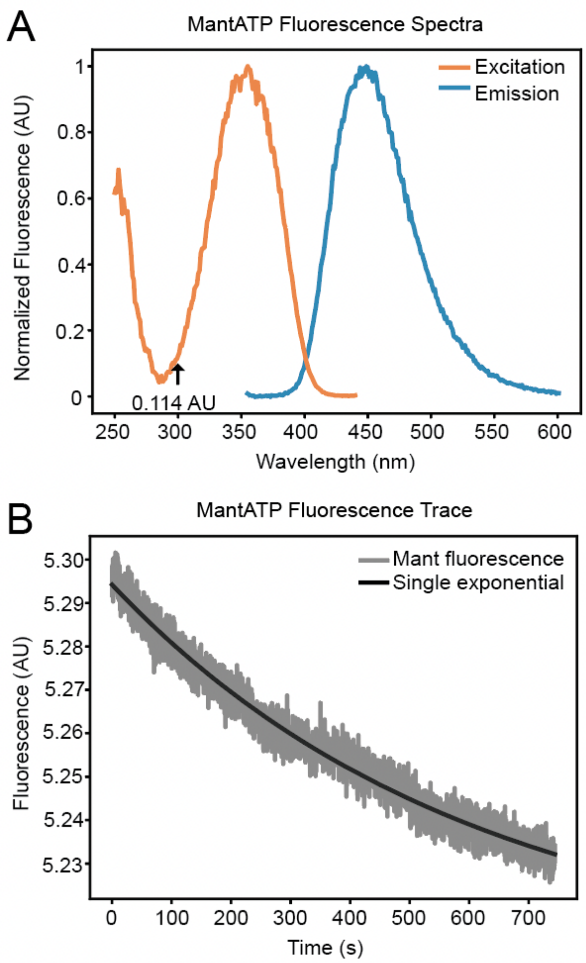


Fig S1: Fluorescent properties of mantATP. A) Excitation (orange) and emission (blue) spectra of mantATP. The active excitation spectrum at 295 nm is 0.114 AU compared to 1 AU at the 350 nm, the excitation peak. B) Raw fluorescence signal of mantATP alone during kinetic acquisition period, with single-exponential fit of fluorescence decrease (presumably due to photobleaching).

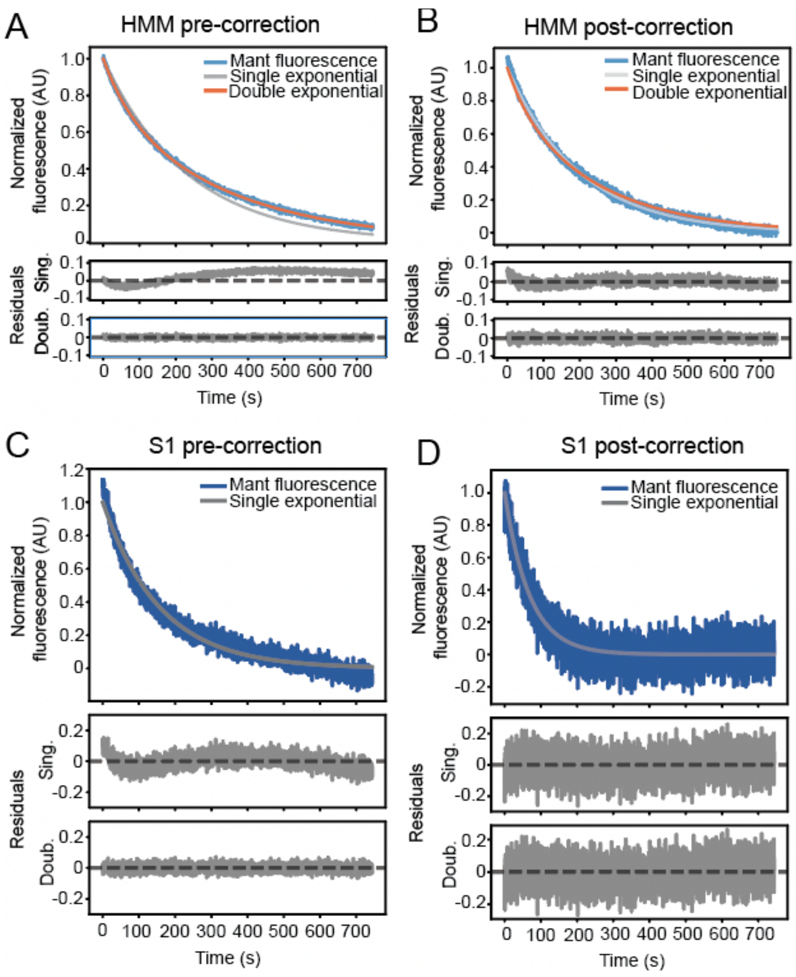


Fig S2: Single exponential fits comparing pre- and post-correction traces of WT-myosin nucleotide turnover. A-B) Sample traces of WT-cHMM nucleotide turnover without correction (A) and with correction for photobleaching (B), both with single and double exponential fits and residuals, showing improvement in single exponential fit upon correcting for mant photobleaching. C-D) Sample traces of WT-cS1 nucleotide turnover pre-correction (C) and post-correction (D) with single and double exponential fits and residuals, demonstrating that photobleaching accounts for double-exponential behavior present in some WT-cS1 traces. Statistical tests indicate that photobleaching corrections eliminate the statistical justification for the second exponential component (Table S1).

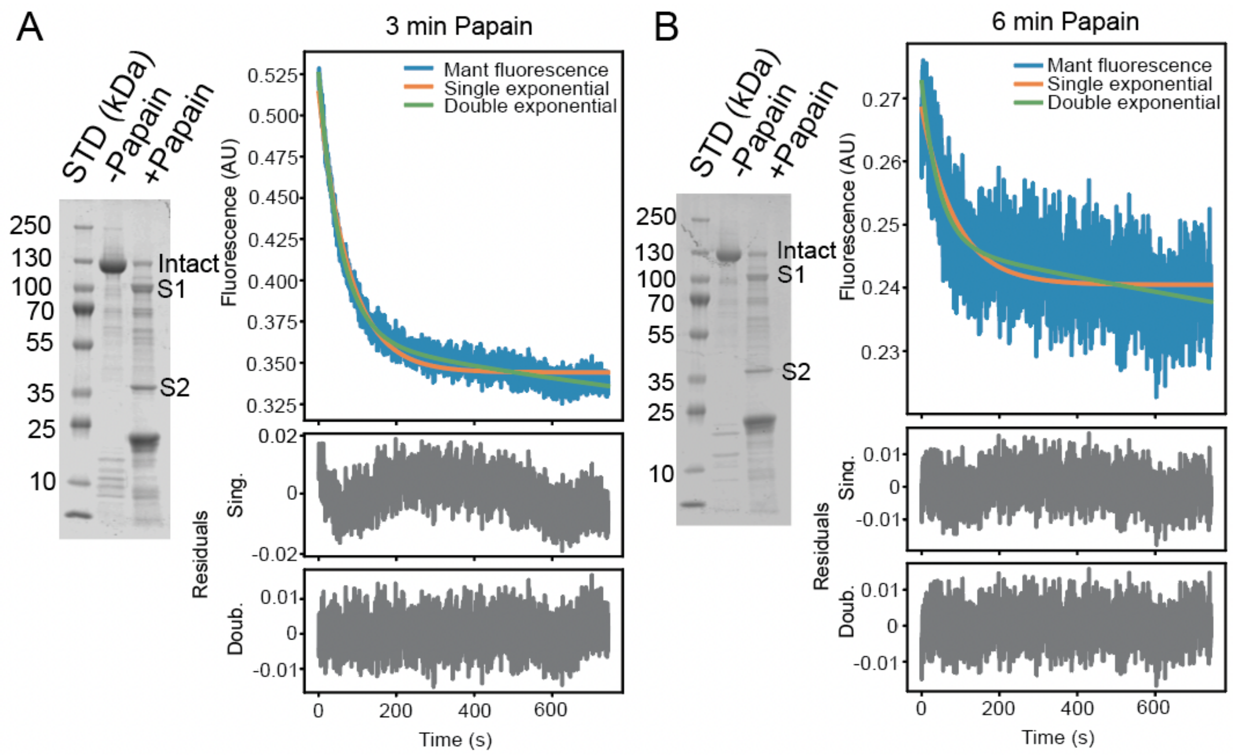


Fig S3: Sensitivity of WT-cHMM digestion to time exposed to papain. A) 3 min papain digestion results in a mixed population with slow and fast populations, likely due to remaining undigested WT-cHMM, visible at 125 kDa in a SDS-PAGE/Coomassie stained gel. B) 6 min papain digestion results in increased-digested samples, wherein the amplitude of the signal (here seen as SNR) is diminished, likely due to increased fragmentation of myosin heads by the protease. The relative amount of myosin digestion was approximated by determining the background corrected Coomassie intensity ratio ($I_{digest}$) of (intact heavy chain + papain)/(intact heavy chain no papain), with $I_{digest}$ = 0.21 for 3 min digestion and $I_{digest}$ = 0.12 for 6 min digestion.

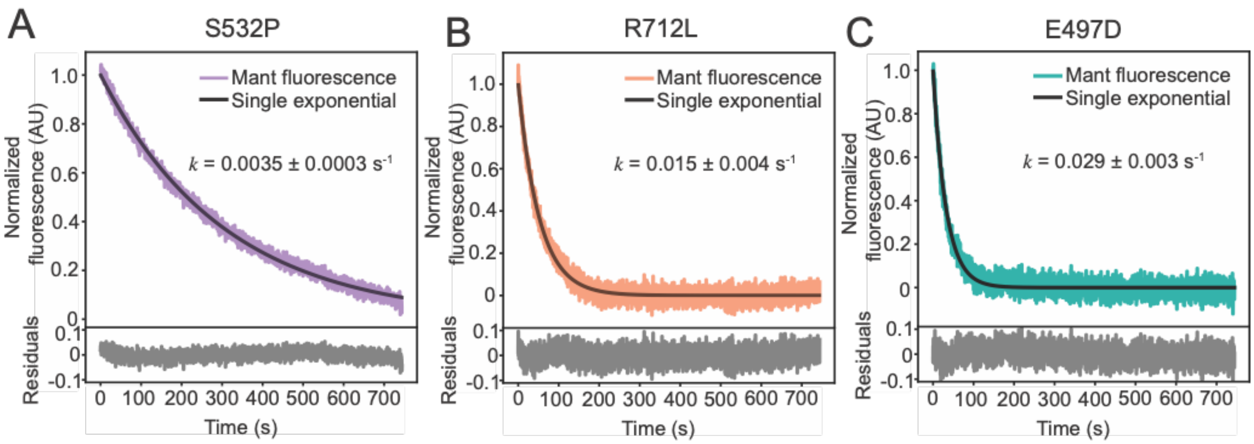


Fig S4: Mutant-cHMM molecules (A: S532P, B: R712L, and C: E497D) turn over nucleotide with single-exponential kinetics, with rates dependent on the mutation.

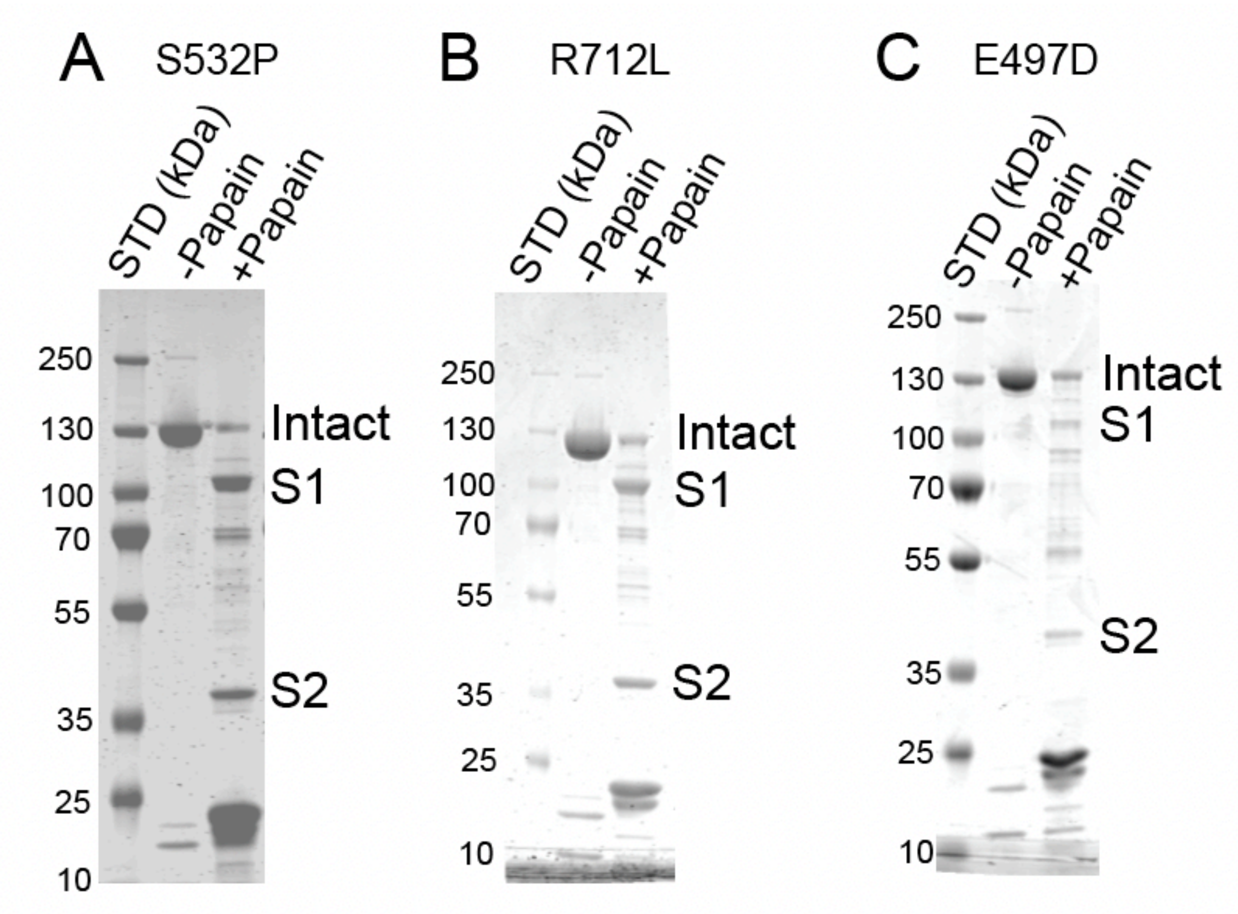


Fig S5: SDS-PAGE/Coomassie stained gel of A) S532P, B) R712L, and C) E497D, showing proteolytic fragmentation by papain.
